# Pathogenic LRRK2 R1441C mutation is associated with striatal alterations

**DOI:** 10.1101/2020.03.11.986455

**Authors:** Harry S. Xenias, Chuyu Chen, Shuo Kang, Suraj Cherian, Xiaolei Situ, Bharanidharan Shanmugasundaram, Giuseppe Scesa, C. Savio Chan, Loukia Parisiadou

**Author notes:** equal contribution (Chuyu Chen, Shuo Kang). Author contributions HSX and SC conducted the electrophysiological measurements. HSX designed and conducted the voltammetric measurements. CC, SK, BS, XS, and GS conducted behavioral testing. HSX and SK conducted statistical analysis. LP, CSC, and HSX wrote the manuscript with input from all co-authors. LP and CSC designed, directed, and supervised the project. All authors reviewed and edited the manuscript.

## Abstract

LRRK2 mutations are associated with both familial and sporadic forms of Parkinson’s disease (PD). Convergent evidence suggests that LRRK2 plays critical roles in regulating striatal function. Here, by using knock-in mouse lines that express the two most common LRRK2 pathogenic mutations—G2019S and R1441C—we investigated how pathogenic LRRK2 mutations altered striatal physiology. We found that R1441C mice displayed a reduced nigrostriatal dopamine release and hypoexcitability in indirect-pathway striatal projection neurons. These alterations were associated with an impaired striatal-dependent motor learning. This deficit in motor learning was rescued following the subchronic administration of the LRRK2 kinase inhibitor Mli-2. In contrast, though a decreased release of dopamine was observed in the G2019S knock-in mice no concomitant cellular and behavioral alterations were found. In summary, our data argue that the impact of LRRK2 mutations cannot be simply generalized. Our findings offer mechanistic insights for devising treatment strategies for PD patients.

## Introduction

The identification of LRRK2 mutations provided important insights into the genetic basis of Parkinson’s disease (PD). Autosomal dominant mutations in LRRK2 are the most common genetic cause of late-onset PD to date ^[1,2]^. Accordingly, compelling evidence from genome-wide association studies has identified *LRRK2* as a risk factor for sporadic PD ^[3-5]^. Patients with LRRK2 mutations exhibit clinical and pathological phenotypes that are indistinguishable from sporadic PD ^[6-8]^, suggesting common disease mechanisms. These findings urge further efforts in understanding mutant LRRK2-induced pathophysiology. The yielded knowledge about the impact of different LRRK2 mutations can be leveraged for therapeutic benefit in PD.

The LRRK2 mutations G2019S and R1441C (hereafter referred to as GS or RC) are commonly found in familial PD ^[9-11]^. These two mutations are located in the kinase and ROC (Ras of complex GTPase) domains of the LRRK2 protein, respectively (**Figure 1a**). It is widely thought that all pathogenic mutations increase LRRK2 kinase activity ^[12, 13]^. Recent studies show that the RC and other related mutations such as R1441G/H result in even greater activation of LRRK2 kinase activity compared to GS mutation ^[1, 13-15]^. Though the literature focuses on the GS mutation, this highlights the importance of determining the impact associated with the RC mutation. The yielded knowledge should, in turn, facilitate the development of therapeutic interventions. Relatedly, previous studies provide support for distinct clinical features associated with different *LRRK2* variants ^[16, 17]^.

**Figure 1.**
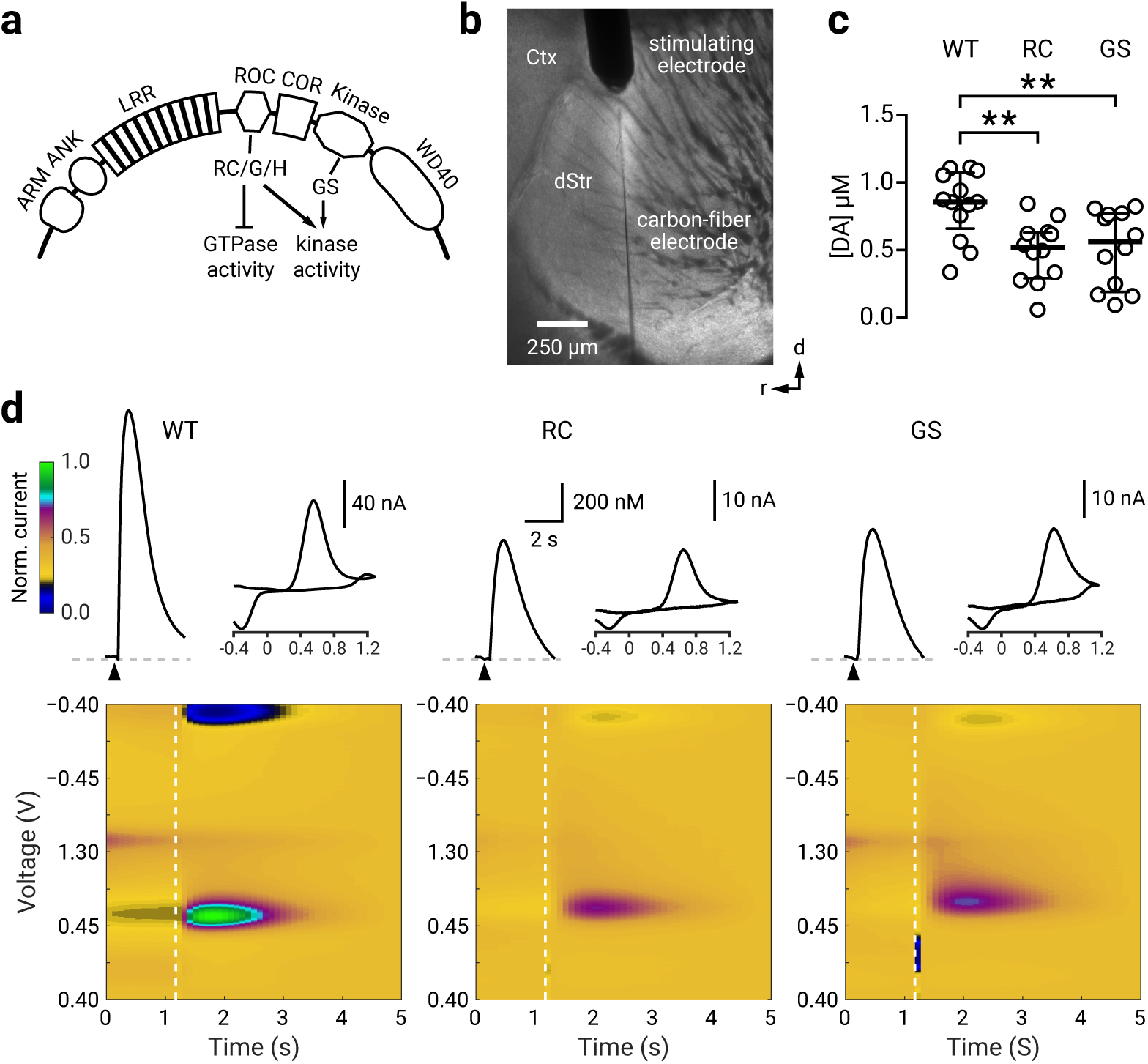
RC andGS LRRK2 mice have decreased nigrostriatal dopamine release. **a**. A schematic diagram of the LRRK2 protein. A family of a Ras-like G-proteins with functionally distinct multi-domains, consisting of armadillo repeats (ARM), ankyrin repeats (ANK), the leucine-rich repeats (LRR), the ROC GTPase domain where the R1441C/G/H mutations reside, and the kinase domain where the G2019S mutation resides. The WD40 domain is involved in membrane binding with ARM and ANK believed to stabilize the electrostatic surfaces of the domains. **b**. Brightfield photomicrograph of a parasagittal slice with a concentric electrical electrode placed into the dorsal striatum; a carbon fiber electrode (CFE) was inserted adjacent to the stimulation site. **c**. Population data showing the peak evoked [DA] in WT (n = 13 recordings), GS (n = 19 recordings), and RC (n = 19 recordings) LRRK2 KI mice. **d**. Representative time courses for [DA] with corresponding cyclic voltammograms and color maps of fast-scanning cyclic voltammetry recordings. Colormaps are normalized to the oxidation current from the WT recording. ** *p* < 0.01

Transgenic mouse models overexpressing LRRK2 GS and RC mutations have hinted at a critical role of LRRK2 in regulating dopamine release in the striatum ^[18, 19]^. However, as the results are highly inconsistent across studies, further efforts are needed to clarify the precise alterations induced by LRRK2 mutations. Here, by using gene-targeted knock-in (KI) mice, we systematically investigated the alterations in striatal electrochemical, electrophysiological, and motor function associated with LRRK2 RC and GS mutations.

## Methods

### Mice

All experiments were in compliance with Northwestern University Animal Care and Use Committee guidelines. For electrophysiological studies, Drd1a-tdTomato mice (Jackson Laboratory 016204) were crossed with R1441C mice (Jackson Laboratory 009346) ^[20]^ and Drd2-eGFP mice (MMRC 000230) were crossed with G2019S mice ^[21]^. All mice were maintained on the C57BL/6 (Jax 000664) background. Heterozygotes for mutant LRRK2 alleles and their littermate controls, or wild-type (WT) mice, were used in all experiments. For electrophysiological recordings, hemizygotes for Drd1a-tdTomato and Drd2-eGFP were used for cellular identification. Mice were group-housed on a standard 12/12 hr light/dark cycle. Both males and females were used in this study.

### Visualized *ex vivo* electrophysiology

Mice at postnatal day 90–110 were anesthetized with a ketamine-xylazine mixture and perfused transcardially with ice-cold aCSF containing the following (in mM): 125 NaCl, 2.5 KCl, 1.25 NaH_2_PO_4_, 2.0 CaCl_2_, 1.0 MgCl_2_, 25 NaHCO_3_, and 12.5 glucose, bubbled continuously with carbogen (95% O_2_ and 5% CO_2_). The brains were rapidly removed, glued to the stage of a vibrating microtome (Leica Instrument), and immersed in ice-cold aCSF. Parasagittal slices containing the dorsal striatum were cut at a thickness of 240 μm and transferred to a holding chamber where they were submerged in aCSF at 37 °C for 30 min and maintained at room temperature thereafter. Slices were then transferred to a small-volume (∼0.5 ml) Delrin recording chamber mounted on a fixed-stage, upright microscope (Olympus). Neurons were visualized using differential interference contrast optics (Olympus), illuminated at 735 nm (Thorlabs), and imaged with a 60× water-immersion objective (Olympus) and a CCD camera (QImaging). Genetically-defined neurons were identified by somatic eGFP or tdTomato fluorescence examined under epifluorescence microscopy with a white (6,500 K) LED (Thorlabs) and appropriate filters (Semrock).

Recordings were made at room temperature (20–22 °C) with patch electrodes fabricated from capillary glass (Sutter Instrument) pulled on a Flaming-Brown puller (Sutter Instrument) and fire-polished with a microforge (Narishige) immediately before use. Pipette resistance was typically ∼3–4 MΩ. For whole-cell current-clamp recordings, the internal solution consisted of the following (in mM): 135 KMeSO_4_, 10 Na_2_phosphocreatine, 5 KCl, 5 EGTA, 5 HEPES, 2 Mg_2_ATP, 0.5 CaCl_2_, and 0.5 Na_3_GTP, with pH adjusted to 7.25–7.30 with KOH. The liquid junction potential for this internal solution was ∼7 mV and was not corrected. For voltage-clamp recordings, neurons were clamped at -80 mV with an internal solution that contained the following (in mM): 125 CsMeSO_3_, 10 Na_2_-phosphocreatine, 5 HEPES, 5 tetraethylammonium chloride, 2 Mg_2_ATP, 1 QX314-Cl, 0.5 Na_3_GTP, 0.25 EGTA, and 0.2% (w/v) biocytin, with pH adjusted to 7.25–7.30 with CsOH. Stimulus generation and data acquisition were performed using an amplifier (Molecular Devices), a digitizer (Molecular Devices), and pClamp (Molecular Devices). For current-clamp recordings, the amplifier bridge circuit was adjusted to compensate for electrode resistance and was subsequently monitored. The signals were filtered at 1 kHz and digitized at 10 kHz. KMeSO_4_ and Na_2_-GTP were from ICN Biomedicals and Roche, respectively. All other reagents were obtained from Sigma-Aldrich.

To determine the excitability of SPNs, the frequency-current (F-I) relationship of each cell was examined with current-clamp recordings. A series of current steps of 500 ms duration were applied beginning at -150 pA and incremented at 25 pA for each consecutive sweep. This protocol was applied until each recorded cell reached maximal firing and after entering a depolarization block. Resting membrane potential was monitored for stability and cells that varied 20% from mean baseline were excluded from the analysis.

Corticostriatal responses were recorded in voltage-clamp as previously described ^[22]^. Electrical stimulation was performed using parallel bipolar tungsten electrodes (FHC) placed in layer 5 of the cortex. Stimulus width and intensity were adjusted via a constant current stimulator (Digitimer), to evoke a first excitatory postsynaptic current (EPSC) with an amplitude of 200–400 pA in the presence of the GABA_A_ receptor antagonist SR95531 (10 µM). Whole-cell access was monitored with a -5 mV pulse throughout the recording. Membrane capacitance (Cm) was determined off-line as Cm = Q_t_ * V_test_, where Qt was calculated as the integral of the transient current elicited by V_test_, a 10 mV voltage step ^[23]^. The paired-pulse ratio (PPR) for a given cell was calculated by taking the average of the ratios of the second EPSC amplitude to the first EPSC amplitude for each recording sweep. Data were excluded if the series resistance of the patch pipette differed by >20% between the two recordings.

### Fast-scanning cyclic voltammetry

Brain slices were prepared as described above in the electrophysiology section. Carbon fiber (7 μm diameter) (Goodfellow) electrodes were fabricated with glass capillary (Sutter) using a puller (Narishige) and fiber tips were hand-cut to 30–100 μm past the capillary tip. The carbon-fiber electrode was held at -0.4 V before each scan. A voltage ramp to and from 1.2 V (400 V/s) was delivered every 100 ms (10 Hz). Before recording, electrodes were conditioned by running the ramp at 60 Hz for 15 min and at 10 Hz for another 15 min and calibrated using 1 μM dopamine hydrochloride (Sigma). Dopamine transients were evoked by electrical stimulation delivered through a concentric, bipolar electrode (FHC) placed in the rostrodorsal striatum (**Figure 1b**) because of its known and important involvement in reward and motivation ^[24-27]^ and motor learning and control ^[27-29]^. A single electrical pulse (300 µA, 0.2 ms) was used ^[30, 31]^. Data were acquired with an amplifier (Molecular Devices), a digitizer (Molecular Devices), and pClamp (Molecular Devices). For each slice, four measurements were made and then averaged. The custom analysis was written in MATLAB (MathWorks). The voltammogram and peak oxidative current amplitudes of the dopamine transient were measured. Experiments were rejected when the evoked current did not have the characteristic electrochemical signature of dopamine.

### Behavioral tests

Motor learning was assessed with an accelerating rotarod in WT, RC, and GS mice. The task started when the mice were around postnatal day 60 using a rotarod apparatus (Panlab) equipped with a mouse rod (3 cm diameter) and set to 4–40 rpm acceleration over 300 s. The task consisted of eighteen daily sessions (five trials per session; intertrial-interval = 15 s, max trial duration = 300 s) divided into two phases ^[32]^ (**Figure 3a**). Specifically, during the dopamine receptor antagonism phase (session 1–5), mice were systemically injected (i.p.) 30 min prior to testing with one of the following: 0.9% saline, cocktail of 1 mg/kg SCH23390 + 1 mg/kg eticlopride, 1 mg/kg SCH23390 or 1 mg/kg eticlopride. Following a 72-hr break, mice were then tested for another thirteen sessions (drug-free recovery phase). To determine if LRRK2 mutation contributes to motor learning deficit, a cohort of RC mice was administered with 5 mg/kg MLi-2 (LRRK2 inhibitor) ^[33, 34]^ 60 min prior to all daily sessions and saline or cocktail of 1 mg/kg SCH23390 + 1 mg/kg eticlopride was given 30 min before the first five daily sessions. All the injections were given i.p (0.1 ml/kg). A separate cohort of WT, RC, and GS mice underwent the accelerating rotarod task for the initial drug-treated phase. Before the initiation of the drug-free recovery phase (after the 72-hr break), mice were assessed individually in 56 x 56 cm open-field arenas in noise-canceling boxes and illuminated by dim red lights. Session (five-minute duration) started when mice were placed in the center of the arena. Locomotor activity was analyzed by the LimeLight (Actimetrics) software and reported as distance traveled.

### Western blot analysis

Mice were treated with either MLi-2 (5 mg/kg, i.p.) or vehicle for 60 min; the striata were dissected and rapidly homogenized in four volumes of ice-cold Buffer A (0.32 M sucrose, 5 mM HEPES, 1 mM MgCl_2_, 0.5 mM CaCl_2_, pH 7.4) supplemented with Halt protease and phosphatase inhibitor cocktail (Thermo Fisher Scientific) using a Teflon homogenizer (12 strokes). Homogenized striatal extract was centrifuged at 1,400 *g* for 10 min. 20 μg of the supernatant were separated by 4–12% NuPage Bis-Tris PAGE (Thermo Fisher Scientific) and transferred to membranes using the iBlot nitrocellulose membrane Blotting system (Thermo Fisher Scientific) by following manufacturer’s protocol. Primary antibodies specific for pS935 LRRK2 (Abcam, ab230261 1:1000), total LRRK2 (Abcam, ab133476), and β-actin (Thermo Fisher Scientific, MA1-045 1:3000) were used. Secondary anti-mouse and anti-rabbit antibodies were from Thermo Fisher Scientific Membranes were incubated with Immobilon ECL Ultra Western HRP Substrate (Millipore) for 3 min prior to image acquisition. Chemiluminescent blots were imaged with iBright CL1000 imaging system (Thermo Fisher Scientific).

### Experimental design and statistical analyses

General graphing and statistical analyses were performed with MATLAB (MathWorks), SAS (SAS Institute), and Prism (GraphPad). Custom analysis codes are available on GitHub (https://github.com/chanlab). Sample size (n value) is defined by the number of observations (i.e., neurons or cells). No statistical method was used to predetermine the sample size. Unless noted otherwise, data are presented as median values ± median absolute deviations as measures of central tendency and statistical dispersion, respectively. Box plots are used for graphic representation of population data ^[35, 36]^. The central line represents the median, the box edges represent the interquartile ranges, and the whiskers represent 10–90^th^ percentiles. Normal distributions of data were not assumed for electrophysiological data. Comparisons for unrelated samples were performed using Mann–Whitney *U* test at a significance level (α) of 0.05. Unless noted otherwise, exact P values are reported. Behavioral data are presented as mean ± standard error of the mean and were analyzed with either two-way or three-way ANOVA with repeated measures followed by Tukey *post hoc* tests.

## Results

### R1441C and G2019S LRRK2 mice have decreased nigrostriatal dopaminergic transmission

Studies that examine dopamine transmission in LRRK2 mutant mice have not yielded consistent results ^[18, 37-40]^. This was in part due to the employment of bacterial artificial chromosomes (BAC) or other transgenic lines confounded by unintended genomic alterations in expression patterns and levels of endogenous LRRK2 ^[41, 42]^. KI mouse models serve as a well-validated approach for studying LRRK2 mutations in relevant cell types at the physiologically-relevant expression level. In this study, we used adult LRRK2 RC and GS KI mice that express mutant LRRK2 proteins under the regulation of the endogenous promoter. We first examined nigrostriatal dopamine transmission with fast-scanning cyclic voltammetry in *ex vivo* striatal tissues from WT, RC, and GS mice (**Figure 1b**). Here, we found a decrease in evoked dopamine release in RC mice compared to WT ([DA]_WT_ = 856.5 ± 198.4 nM, n = 13 recordings; [DA]_RC_ = 516.7 ± 148.6 nM, n = 19 recordings; *p =* 0.0024, Mann-Whitney *U* test). In addition, we found a similar decrease in evoked dopamine release in GS mice compared to WT ([DA]_GS_ = 562.4 ± 228.3 nM, n = 19 recordings; *p =* 0.0030, Mann-Whitney *U* test). There were no differences in evoked dopamine release between RC and GS mice (*p =* 0.89, Mann-Whitney *U* test) (**Figure 1c & d**).

### iSPNs have decreased excitability in R1441C LRRK2 mice

It is established that dopamine modulates the excitability of striatal projection neurons in naive animals ^[43-47]^. Accordingly, the disruption of dopamine signaling profoundly alters the intrinsic properties of SPNs ^[22, 48]^. Given our finding that both RC and GS mice had decreased dopamine release, we sought to examine if direct- and indirect-pathway striatal projection neurons (dSPNs and iSPNs) have altered excitability in the LRRK2 mutant mice. Whole-cell, current-clamp recordings were performed on identified SPNs in WT, RC, and GS mice (**Figure 2**).

**Figure 2.**
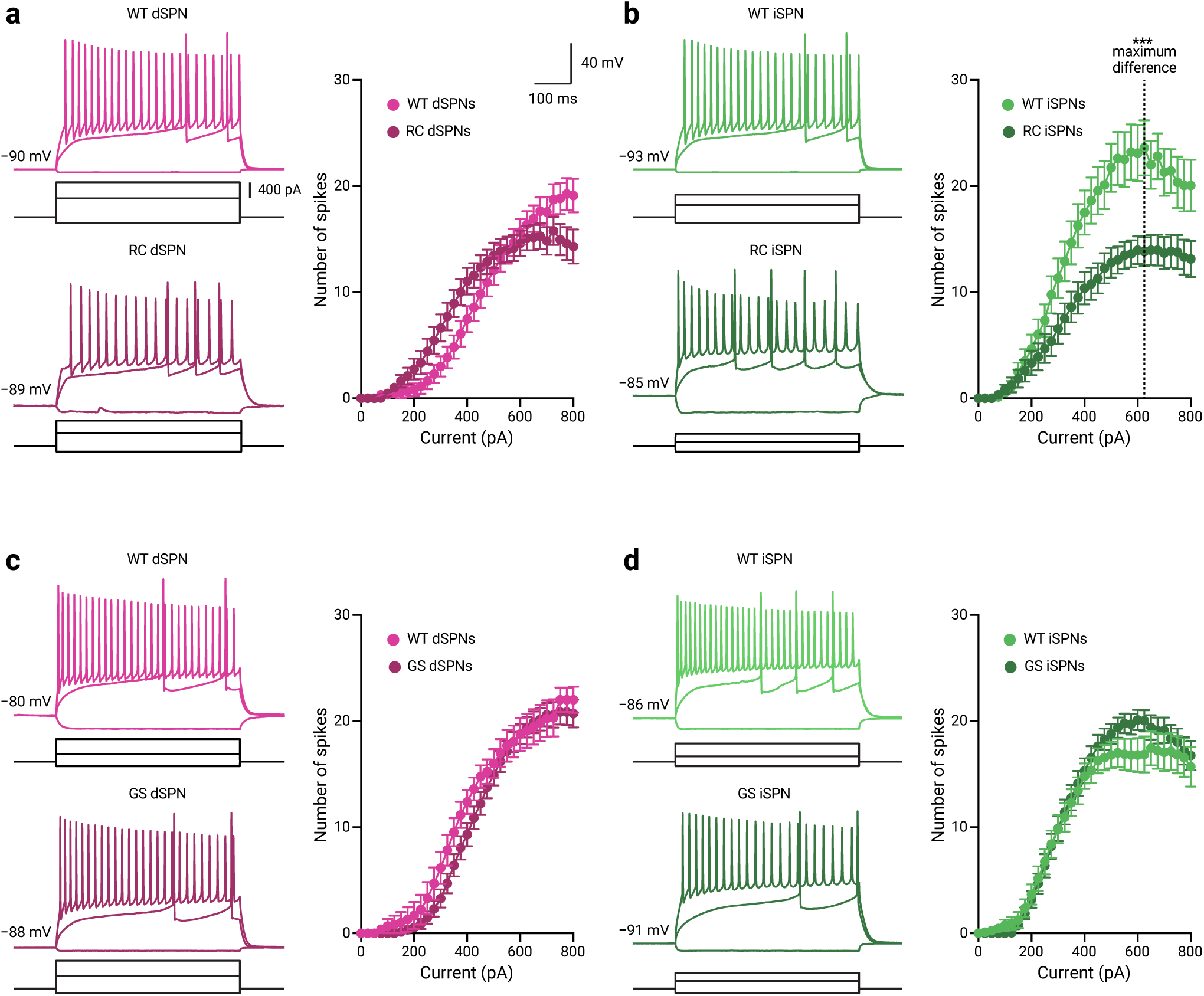
iSPNs in RC LRRK2 mice have decreased excitability. **a–b**. Representative whole-cell current-clamp recordings of dSPNs and iSPNs in WT and RC LRRK2 KI mice. The three injection steps shown elicited the voltage responses to first hyperpolarizing current, the first occurrence of action potentials, and the maximum firing, respectively. The half-maximum firing values in the population data of WT mice for dSPNs (I = 475 pA, n = 20 recordings) and iSPNs (I = 325 pA, n = 20 recordings) were used for statistical comparison with the population F-I data of RC LRRK2 KI mice for dSPNs (n = 30 recordings) and Ispns (n = 23 recordings). The dashed line represents the current where the maximal difference between the F-I functions for the iSPNs in WT and RC mice was calculated (I = 625 pA). **c–d**. Representative whole-cell current-clamp recordings and population data for SPNs of WT and GS LRRK2 KI mice. The half-maximum firing values in the population data ofWT for dSPNs (I = 375 pA, n = 13 recordings) and iSPNs (I = 325 pA, n = 20 recordings) were used for statistical comparison with the population F-I data of GS LRRK2 KI mice for dSPNs (n = 29 recordings) and iSPNs (n = 28 recordings). F-I data are shown as mean ± standard error of the mean. *** *p* < 0.001

**Figure 3.**
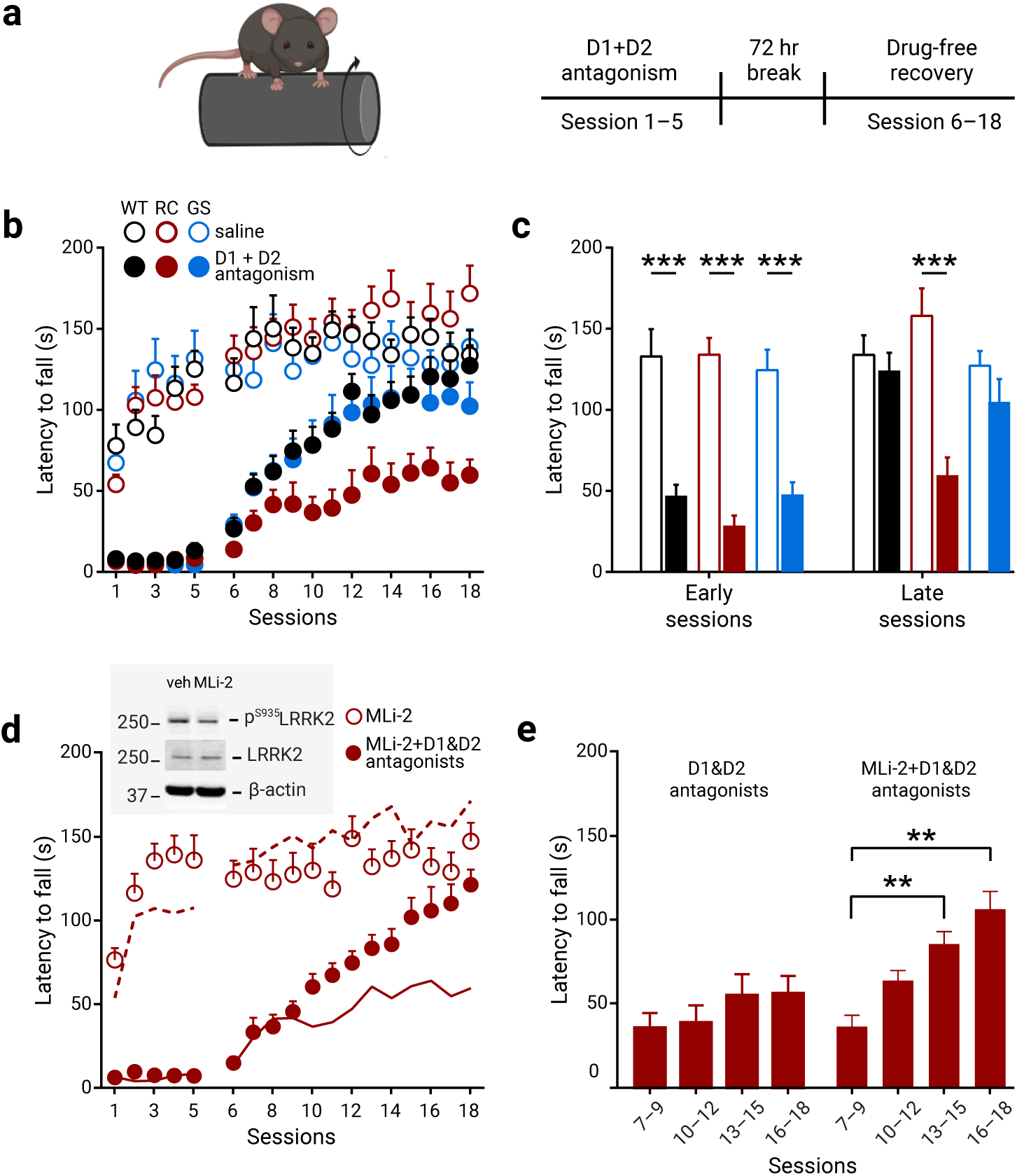
RC KI mice show dopamine-dependent motor learning impairments. **a**. Schematic of the rotarod training paradigm. Mice were assessed over a total of 18 daily sessions where every daily session consisted of five trials. WT, GS, and RC mice were administered either saline or a cocktail of D1 receptor (SCH23390) and D2 receptor (eticlopride), 1 mg/kg of each antagonist 30 min prior to daily sessions, and trained for five successive days in an accelerated rotarod. After a 72 h break, the mice were returned to the rotarod for an additional 13 days of a drug-free recovery phase. **b**. Genotype-dependent effect of blocking both D1 and D2 receptors, during the first five sessions on rotarod performance of WT, RC, and GS mice. The average latency in the early (session 6–8) and late (session 16–18) stage in the drug-free recovery phase in **b** is summarized in **c**. Open and filled bars represent the average performance of saline control and drug-treated mice, respectively (saline treated groups: n_WT_ = 10 mice, n_RC_ = 10 mice, and n_GS_ = 9 mice; D1 + D2 receptor antagonist treated groups: n_WT_ = 13 mice, n_RC_ = 11 mice, and n_GS_ = 11 mice; *** *p <* 0.001 vs genotype-matched saline control). **d**. The motor learning deficit induced by dopamine antagonism was not observed in RC mice pre-treated with the LRRK2 inhibitor MLi-2, which was administered daily, 60 min prior to training. group of RC mice that were then treated with either saline (open circles) or antagonist cocktail (closed circles) for the first 5 sessions. Dash and solid lines respectively represent the saline control or dopamine receptor antagonist treated groups. Inset shows a decrease in S935 LRRK2 phosphorylation, which reflects LRRK2 kinase activity in WT mice after 60 min Mli-2 administration. **e**. Average latency in blocks of 3 sessions during the drug free phase from **d**. MLi-2 with saline: n = 12 mice; MLi-2 with D1+D2 antagonists: n = 11 mice; ** *p* < 0.01).

We found no differences in the number of spikes in dSPNs between WT and RC mice for the current (I) that elicited the half-maximum firing from dPSNs in the WT mice (I = 475 pA, output_WT_ = 12.0 ± 4.0 spikes, n = 20 recordings; output_RC_ = 14.0 ± 3.0 spikes, n = 30 recordings; *p =* 0.21, Mann–Whitney *U* test) (**Figure 2a**). In contrast, there was a decrease in the number of evoked spikes for the iSPNs in RC mice for the current that yielded the half-maximum firing from iSPNs in the WT mice (I = 325 pA, output_WT_ = 14.0 ± 5.0 spikes, n = 20 recordings; output_RC_ = 6.0 ± 6.0 spikes, n = 23 recordings; *p =* 0.037; Mann–Whitney *U* test) (**Figure 2b**). To quantify the decreased excitability in the iSPNs of RC mice, we compared input-output functions for the WT and RC mice. As shown in **Figure 2b**, the maximal difference in the spike output was observed at I = 625 pA; a lower number of spikes were elicited in iSPNs of RC mice compared to WT mice (output_WT_ = 22.0 ± 4.5 spikes, n = 17 recordings; output_RC_ = 15.0 ± 5.0 spikes, n = 23 recordings; *p =* 0.0010, Mann-Whitney *U* test) (**Figure 2b**). In contrast, we found no detectable differences in GS mice as measured with the half-maximum firing responses for either dSPNs (output_WT_ = 8.0 ± 5.0 spikes, n = 13 recordings; output_GS_ = 11.0 ± 3.0 spikes, n = 29 recordings; *p* = 0.12, Mann-Whitney *U* test) or iSPNs (output_WT_ = 10.5 ± 4.8 spikes, n = 20 recordings; output_GS_ = 12.0 ± 4.0 spikes, n = 28 recordings, *p* = 0.73, Mann-Whitney *U* test) (**Figure 2c & d**).

As the excitability of SPNs is a function of dendritic structure ^[49]^, we asked whether an increase in membrane capacitance (Cm) could account for the decreased excitability seen in the iSPNs of RC mice. We found an increased membrane capacitance for iSPNs in RC mice compared to WT mice (Cm_WT_ = 68.8 ± 13.9 pF, n = 27; Cm_RC_ = 85.2 ± 13.3 pF, n = 27; *p* = 0.039, Mann-Whitney *U* Test). In contrast, no change in membrane capacitance was observed for dSPNs between RC and WT mice (Cm_WT_ = 79.4 ± 19.5 pF, n = 25 recordings; Cm_RC_ = 85.1 ± 16.0 pF, n = 38 recordings; *p* = 0.98, Mann-Whitney *U* Test) (**Figure 2–figure supplement 1**). We also examined for changes in capacitance in GS mice. In line with previous findings ^[50, 51]^, we found no changes in the intrinsic properties of SPNs in the GS mice. We found no differences in membrane capacitance between GS mutant or WT mice for dSPNs (Cm_WT_ = 88.1 ± 10.8 pF, n = 15 recordings; Cm_GS_ = 95.4 ± 12.0 pF, n = 27 recordings; *p* = 0.29, Mann-Whitney *U* Test) or iSPNs (Cm_WT_ = 74.6 ± 12.3 pF, n = 19 recordings; Cm_GS_ = 81.8 ± 15.2 pF, n = 30 recordings; *p* = 0.51, Mann-Whitney *U* Test) (**Figure 2–figure supplement 1**).

The hypoexcitability of iSPNs in models of PD has been suggested as a homeostatic response to an increased corticostriatal transmission ^[50, 52, 53]^, we thus examined possible presynaptic changes in corticostriatal transmission by measuring the paired-pulse ratios (PPRs) of the corticostriatal responses in WT and RC mice (see Methods). There were no differences in the PPRs for either dSPNs (PPR_WT_ = 1.31 ± 0.10, n = 14 recordings; PPR_RC_ = 1.33 ± 0.15, n =17 recordings; *p =* 0.18, Mann-Whitney *U* test) or iSPNs (PPR_WT_ = 1.28 ± 0.14, n = 13 recordings; PPR_RC_ = 1.54 ± 0.25, n = 18 recordings; *p =* 0.11, Mann-Whitney *U* test) (**Figure 2–figure supplement 2**). Unexpectedly, we found an increase in the 95%–5% EPSC decay time of iSPNs in RC mice compared to WT mice (decay_WT_ = 61.1 ± 38.7 ms, n = 13 recordings; decay_RC_ = 139.5 ± 81.1 ms, n = 18 recordings; *p* = 0.037, Mann-Whitney *U* test) (**Figure 2–figure supplement 2**). In contrast, there was no difference in EPSC decay times for dSPNs between WT mice and RC mice (decay_WT_ = 88.7 ± 40.3 ms, n = 14 recordings; decay_RC_ = 182.9 ± 136.0 ms, n = 17 recordings; *p =* 0.13, Mann-Whitney *U* test) (**Figure 2–figure supplement 2**).

### R1441C mice show impaired dopamine-dependent motor learning

The dorsal striatum plays a critical role in habit and motor learning ^[54-57]^. Given the reduction of dopamine release in the dorsal striatum in RC and GS KI mice (**Figure 1**) we sought to investigate how a disruption to dopamine signaling would affect motor learning. Specifically, we first systemically injected D1 and D2 dopamine receptor antagonists (SCHSCH23390 and eticlopride, referred to as “antagonists cocktail”) to determine the roles of D1 and D2 dopamine receptor-mediated signaling on motor learning in RC and GS mice. We then used an accelerating rotarod task—a well-established paradigm for assessing striatally dependent motor learning ^[32, 58-60]^.

Consistent with previous studies that RC KI mice display no overt abnormalities in striatum-dependent motor learning under basal conditions ^[20, 61]^, we found that saline treated RC mice learned equally well on the rotarod task (**Figure 3b**). Our data consistent with prior reports in WT mice ^[32, 58]^ showed that mice treated with antagonists cocktail during the initial five-day training period (session 1–5; **Figure 3a**) have dramatic impairments in the rotarod performance, regardless of genotype.

Here, we showed that upon 72 hrs of recovery from the last antagonist administration, the latency to fall for WT, RC, and GS mice although initially degraded (compared to saline treated controls), (**Figure 3b**), increased over subsequent sessions in different manners (saline: n_WT_ = 10 mice, n_RC_ = 10 mice, and n_GS_ = 9 mice; D1 + D2 receptor antagonists: n_WT_ = 13 mice, n_RC_ = 11 mice, and n_GS_ = 11 mice; treatment x genotype x session interaction, F_102, 1887_ = 1.95, *p <* 0.001, 3-way ANOVA with repeated measures) (**Figure 3b**). The average latency to fall in the drug-treated WT and GS mice showed no difference compared to saline-treated controls over the training sessions (session 16–18; *p =* 1.0, latency_WT_saline_ = 137.2 ± 11.9 s, latency_WT_antagonists_ = 117.2 ± 8.2 s; *p =* 0.98, latency_GS_saline_= 130.9 ± 8.7 s, latency_GS_antagonists_= 99.2 ± 15.6 s, 3-way ANOVA, repeated measures) (**Figure 3b & c, Figure 3–table supplement 1**). In contrast, dopamine receptor antagonism unmasked a deficit in RC KI mice; they failed to reach the performance of their corresponding saline-treated controls over the training sessions (trial x genotype x drug interaction: F_6,116_ = 2.24, *p* = 0.044; latency_RC_saline_ = 162.4 ± 17.0 s, latency_RC_antagonists_ = 58.5 ± 11.1 s, *p* < 0.001, 3-way ANOVA with Tukey *post hoc* test) (**Figure 3c**). Similarly, RC KI mice failed to reach the performance of their dopamine D1 and D2 receptor antagonist treated WT controls (trial x genotype x drug interaction: F_6,116_ = 2.24, *p* = 0.044; latency_WT_antagonists_ vs latency_RC_antagonists_, *p* < 0.043, 3-way ANOVA with Tukey *post hoc* test) (**Figure 3c**).

Given that LRRK2 mutations stimulate LRRK2 kinase activity, we examined whether increased kinase activity in RC mice contributes to this motor learning deficit. Therefore, we treated RC mice with the LRRK2 inhibitor MLi-2 ^[33]^ prior to D1 and D2 receptor antagonism. We found motor improvement across behavioral sessions during the drug-free phase of MLi-2-treated mice; this was evident with an increased latency to fall across successive trials (time x treatment interaction, F_17, 374_ = 12.45, *p* < 0.001, two-way ANOVA) (**Figure 3d**). This inference was confirmed by analyzing the average latency in mice with and without MLi-2 treatment. Increase in latency was only observed following MLi-2 injections (time x treatment interaction, F_3, 60_ = 10.98, *p* < 0.001; for MLi-2 with D1+D2 antagonism: *p* = 0.002, latency_session_7–9_ = 38.7 ± 7.0 s, latency_session_10–12_ = 67.7 ± 6.4 s; *p* = 0.001, vs latency_session_13-15_ = 90.6 ± 7.9 s; *p* < 0.001, vs latency_session_16–18_ = 112.8 ± 11.0 s, two-way ANOVA with Tukey *post hoc* tests) (**Figure 3e**). The pre-treatment of MLi-2 decreased LRRK2 S935 phosphorylation—a well-characterized readout of LRRK2 kinase activity ^[62, 63]^—in striatal extracts (**Figure 3d**). In summary, the accelerated rotarod paradigm with dopamine antagonist treatment unmasked the kinase-mediated deficits of striatal motor learning in RC mice.

To rule out the possibility that the impaired learning of RC mice was a lingering effect of the antagonist treatment, a subgroup of RC mice that received the administration of dopamine receptor antagonists were returned to their home cages without rotarod training (here referred to as “untrained mice”). After the 72-hour break, these mice were tested on the rotarod task; their performance was improved compared to the RC mice that underwent training in the rotarod task immediately following systemic administration of dopamine receptor antagonists, as measured by latency to fall (main effect of treatment, F_1, 260_ = 189.2, *p* < 0.0001, two-way ANOVA) (**Figure 3–figure supplement 1**).

To examine the contributions of dopamine receptor subtype in shaping motor learning on the rotarod task, we administered D1 and D2 receptor antagonists separately during the initial acquisition. The administration of SCH23390 or eticlopride impaired the performance during the initial phase of the rotarod training across all genotypes (session 1–5), compared to their saline-treated controls (**Figure 3–figure supplement 1**. While D1 receptor antagonism led to an immediate improvement in all genotypes in the drug-free phase (**Figure 3–figure supplement 1**), D2 receptor antagonism resulted in a delayed improvement in performance (**Figure 3–figure supplement 1**; **Figure 3–table supplement 1**). In addition, WT and GS mice treated with D2 receptor antagonist reached the performance of their saline controls by the end of the task (latency_GS_eticlopride_ = 105.9 ± 9.2 s, latency_GS_saline_, = 130.9 ± 8.7 s; *p* = 1.0, latency_WT_eticlopride_ = 117.8 ± 22.4 s, latency_WT_saline_ = 137.2 ± 11.9 s; *p* = 0.9899, 3-way ANOVA, Tukey *post hoc* test). In contrast, RC mice did not exhibit the performance of their saline controls across sessions (latency_RC_eticlopride_ = 106.3 ± 12.7 s, latency_RC_saline_, = 162.4 ± 17.0 s *p* = 0.016; 3-way ANOVA with Tukey *post hoc* test).

To show that the motor learning impairment in RC mice were not confounded by locomotor deficits, we examined the locomotor behavior of mice in the open field arena. WT, RC, and GS mice that underwent dopamine antagonism and an initial five-day rotarod training were assessed by the total distance traveled. We found no differences between any of the groups except for a significant main effect of drug treatment (distance_WT_saline_ = 28.48 ± 3.36 m; distance_WT_antagonists_ = 24.44 ± 2.90 m; distance_RC_saline_ = 36.02 ± 2.26 m; distance_RC_antagonists_ = 30.62 ± 4.48 m; distance_GS_saline_ = 31.91 ± 2.19 m; distance_GS_antagonists_ = 30.85 ± 3.49 m; treatment effect, F_3,83_ = 4.63, *p* = 0.0048; genotype effect, F_2,83_ = 0.61, *p* = 0.54; treatment x genotype effect, F_6,83_ = 1.94, *p* = 0.084, 3-way ANOVA) (**Figure 3–figure supplement 1**). In summary, disruption of dopamine signaling during rotatord training task specifically interferes with striatal motor learning in the RC mice, which otherwise exhibit no deficits in naturalistic behavior such as locomotion.

## Discussion

While a number of transgenic animal models have been generated to interrogate dysfunction associated with mutant LRRK2, the findings so far have been inconsistent. The most parsimonious explanation is the difference in expression levels of mutant LRRK2 in the presence of endogenous LRRK2, as well as the different promoters used to drive the expression of mutant protein ^[18]^. In this study, by using RC and GS KI mice that express the mutant LRRK2 protein with endogenous expression patterns and levels, we performed a systematic comparative study between these two Lrrk2 KI mouse models to resolve conflicting reports of dopamine transmission ^[21, 51]^. We found a decrease in nigrostriatal dopamine release in both RC and GS mice. Additionally, iSPNs in the RC mice showed a selective decrease in excitability, while no differences were found in dSPNs. The alterations in the cellular properties of iSPNs in RC mice were paralleled with impairments in dopamine-dependent striatal motor learning. We currently do not fully understand the dissociation between impaired dopamine release and striatal alterations. We speculate that PKA signaling at the striatal level may be dysregulated due to RC mutation ^[64, 65]^. This is in line with our finding that increased synaptic PKA activities are observed only in RC and not GS striatal synaptic fractions ^[15,66]^

To our knowledge, this study is the first to examine evoked dopamine release in the RC KI mice. Our results are at odds with an earlier report that measured basal dopamine content in RC KI mice and found no changes using bulk tissue HPLC, which lacks the spatial and temporal resolution that fast-scanning cyclic voltammetry offers ^[20]^. On the other hand, our data corroborate the finding that stimulated catecholamine release in cultured chromaffin cells of RC mice had a 50% reduction of dopamine release, ^[20]^. The decrease of evoked dopamine release could be attributed to the role of LRRK2 in regulating presynaptic vesicle release ^[67-70]^. In particular, LRRK2 targets downstream Rab proteins that are involved in vesicular trafficking to the plasma membrane and direct neurotransmitter release ^[12, 71]^. Specifically, we recently demonstrated that RC mutation resulted in higher phosphorylation of Rab8A—a downstream LRRK2 substrate—in synaptic striatal extracts, compared to GS mutation ^[15]^. In addition, the LRRK2 RC mutation leads to a synaptic translocation and dysregulation of excitatory synaptogenesis and transmission via an increased PKA-mediated regulation of GluR1 in SPNs ^[66]^. It remains to be determined if these alterations are specific to iSPNs and if they are associated with the hypoexcitability observed in our study. Overall, our results show that the RC mutation has a more profound effect on striatal physiology.

In the present study we showed that GS mutation caused decreased evoked dopamine release. Previous studies that measured dopamine in GS KI mice have been contradictory with one another. Using microdialysis, reduced extracellular levels of dopamine at twelve months but not six months were reported ^[21]^. In contrast, by using amperometry Tozzi et al. reported reduced striatal dopamine levels in mice at six months ^[51]^. Moreover, a recent report showed no difference on peak dopamine dopamine release in slices of three months ^[50]^. However, when ∼70% maximum dopamine release was evoked by a single stimuli, they reported a significant increase in dopamine release in GS mice. There are several possibilities for the variability across the results including age of animals, genetic backgrounds, methods to evaluate dopamine content, and heterogeneity across striatal subregions of dopamine release in different experiments.

Given the importance of nigrostriatal dopamine signaling in striatal motor learning, we assessed motor learning in both RC and GS KI lines using the accelerated rotarod. Consistent with previous studies, we observed no abnormalities in motor learning in LRRK2 mutants under basal conditions ^[20, 61]^. While our data are in agreement with previous studies showing that WT mice that underwent dopamine receptor antagonist treatment have an initial delayed motor performance, which gradually improves over sessions ^[32, 72]^, we found that RC mice failed to improve their performance with time. This was not the case for the GS mice. Notably, our findings argue that increased LRRK2 kinase activity underlies this impairment as pretreatment of the mice with the LRRK2 kinase inhibitor improved the performance of RC mice throughout sessions. It is not clear at this point why RC and not GS mice demonstrate this LRRK2 kinase-dependent motor impairment but this finding is in alignment with emerging data demonstrating that mutations outside of the kinase domain lead to greater increase in kinase activity than those found in the kinase domain itself ^[1]^.

Previously, it was suggested that the combination of dopamine antagonists and rotarod training leads to an experience-dependent learning impairment that involves corticostriatal plasticity ^[32, 73, 74]^. Our data are of particular importance as they mirror subclinical dopaminergic dysfunction and corticostriatal alterations of the asymptomatic LRRK2 mutation carriers ^[75-77]^; our study argues that the behavioral deficit associated with LRRK2 mutation is reversed by LRRK2 targeted therapeutic innervations.

Overall, the investigation of the preclinical symptoms of dopamine and striatal dysfunctions in the mutant LRRK2 KI mice provides the framework for implementing neuroprotective therapies and developing biomarkers to detect and monitor disease progression related to LRRK2 mutations. Specifically, given the striatal alterations we observed were in the RC mutant mice, our study highlights the importance of future studies focused on the mutations of the GTPase domain and its downstream signaling targets for the development of signaling-specific neuroprotective therapies.

## Acknowledgments

This work was supported by Michael J. Fox Foundation for Parkinson’s Research (LP), NIH R01 NS097901 (LP), R01 NS097901 (CSC), R01 NS069777 (CSC), P50 NS047085 (CSC), R01 MH112768 (CSC), R01 MH109466 (CSC), R01 NS088528 (CSC), T32 NS041234 (HSX), F32 NS098793 (HSX). We thank Brianna Berceau for colony management and technical support. We also thank Dr. Heather Melrose for providing the LRRK2 G2019S knock-in mice.

## Figure Legends

**Figure 2−figure supplement 1.**
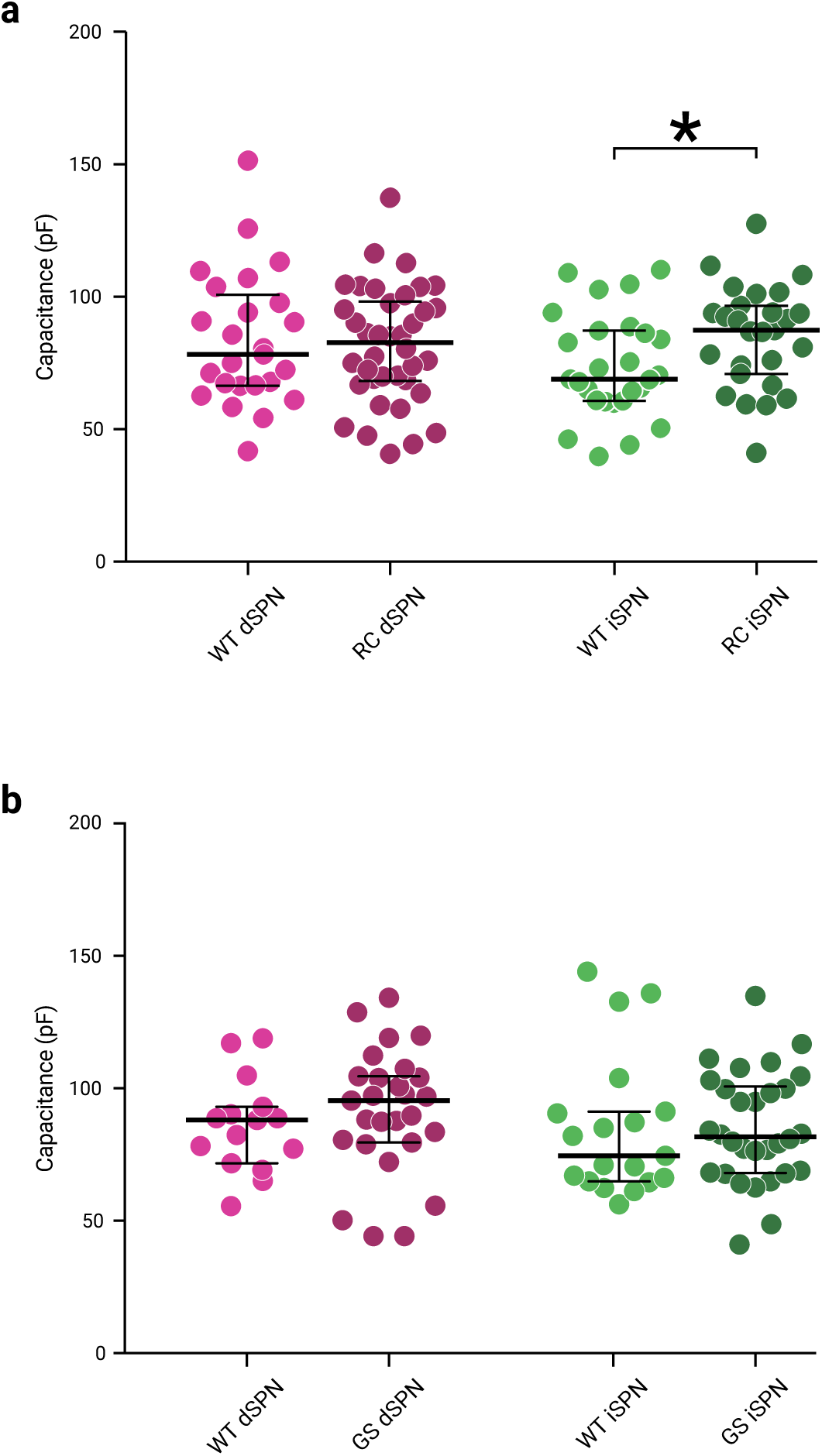
iSPNs have increased membrane capacitance in RC LRRK2 mice. **a**. Scatter plots of membrane capacitance with medians (center line) and interquartile ranges (whiskers) for WT dSPNS (n = 25 recordings) and iSPNS (n = 27 recordings) and RC LRRK2 KI dSPNs (n = 38 recordings) and iSPNs (n = 29 recordings). **b**. Scatter plots of membrane capacitance for SPNs in both WT and GS LRRK2 KI mice.

**Figure 2−figure supplementary 2.**
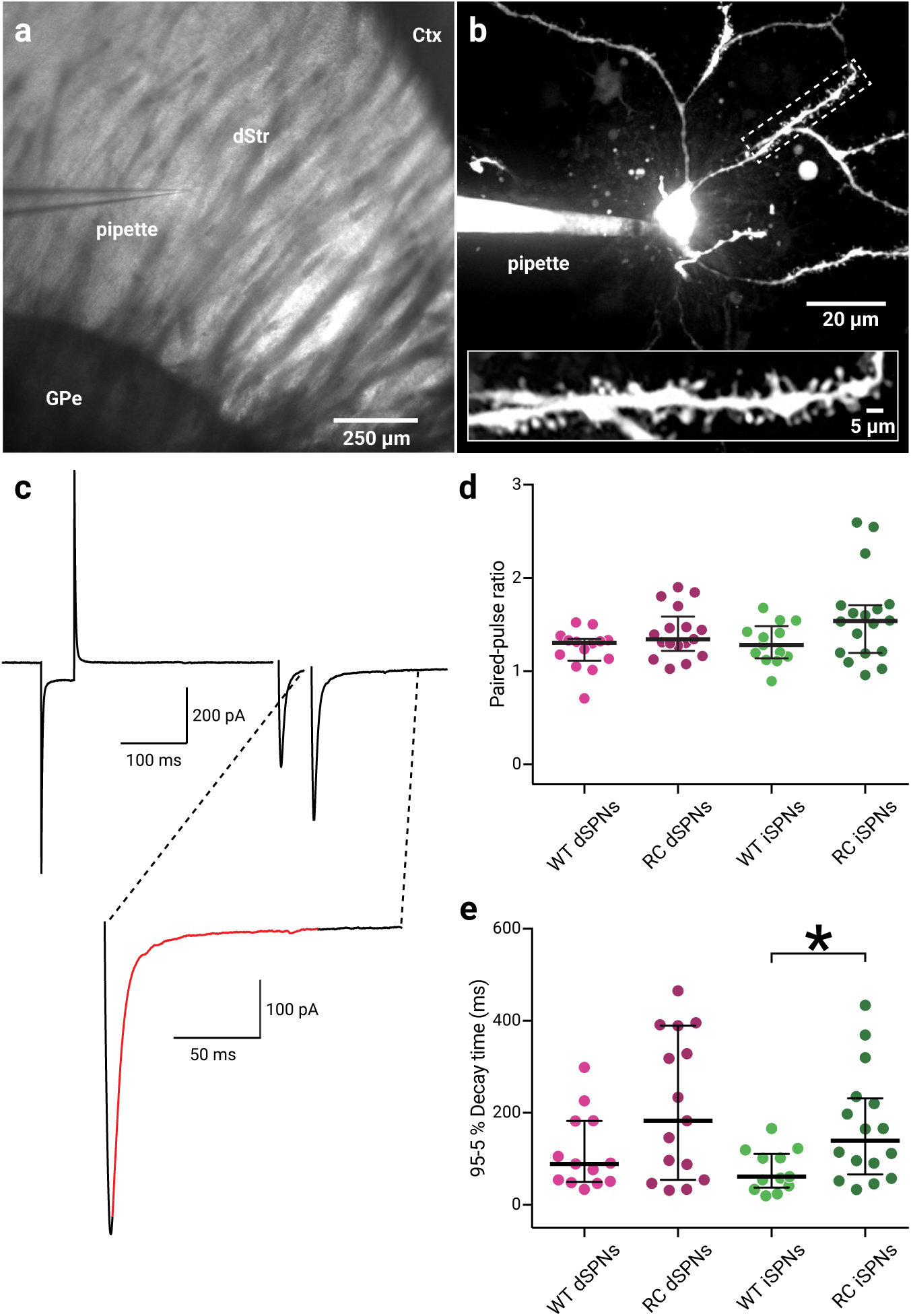
iSPNs have an increased EPSC decay time in RC LRRK2 mice. **a**. Representative brightfield photomicrograph of a WT dSPN in a parasagittal slice voltage-clamped at -80 mV. **b**. Spinning-disk confocal photomicrograph of the same dSPN in the panel to the left. Alexa Fluor 647 was loaded into the cell (via the patch pipette) for visualization. SPNs were identified by morphology and a large number of dendritic spines. Inset: magnified view of the boxed region showing a dendritic segment that is heavily decorated with spines. **c**. *Top left*: Representative voltage-clamp recording of the dSPN shown in **b** above. Paired-pulse stimulation of the cortex at 20 Hz elicited a pair of excitatory postsynaptic currents (EPSC). *Bottom left*: Magnified view of the decay of the second EPSC shown in the boxed region above. The red portion depicts the 95%–5% decay time. **d**. Population data of paired-pulse ratios of WT dSPNs (n = 14 recordings) and iSPNs (n = 13 recordings) and RC dSPNs (n = 17 recordings) and iSPNs (n = 18 recordings). **e**. Population data of the 95%–5% decay times of SPNs from age-matched WT dSPNs (n = 14 recordings) iSPNs (n = 13 recordings) and RC dSPNs (n = 13 recordings) iSPNs (n = 18 recordings).

**Figure 3−figure supplement 1.**
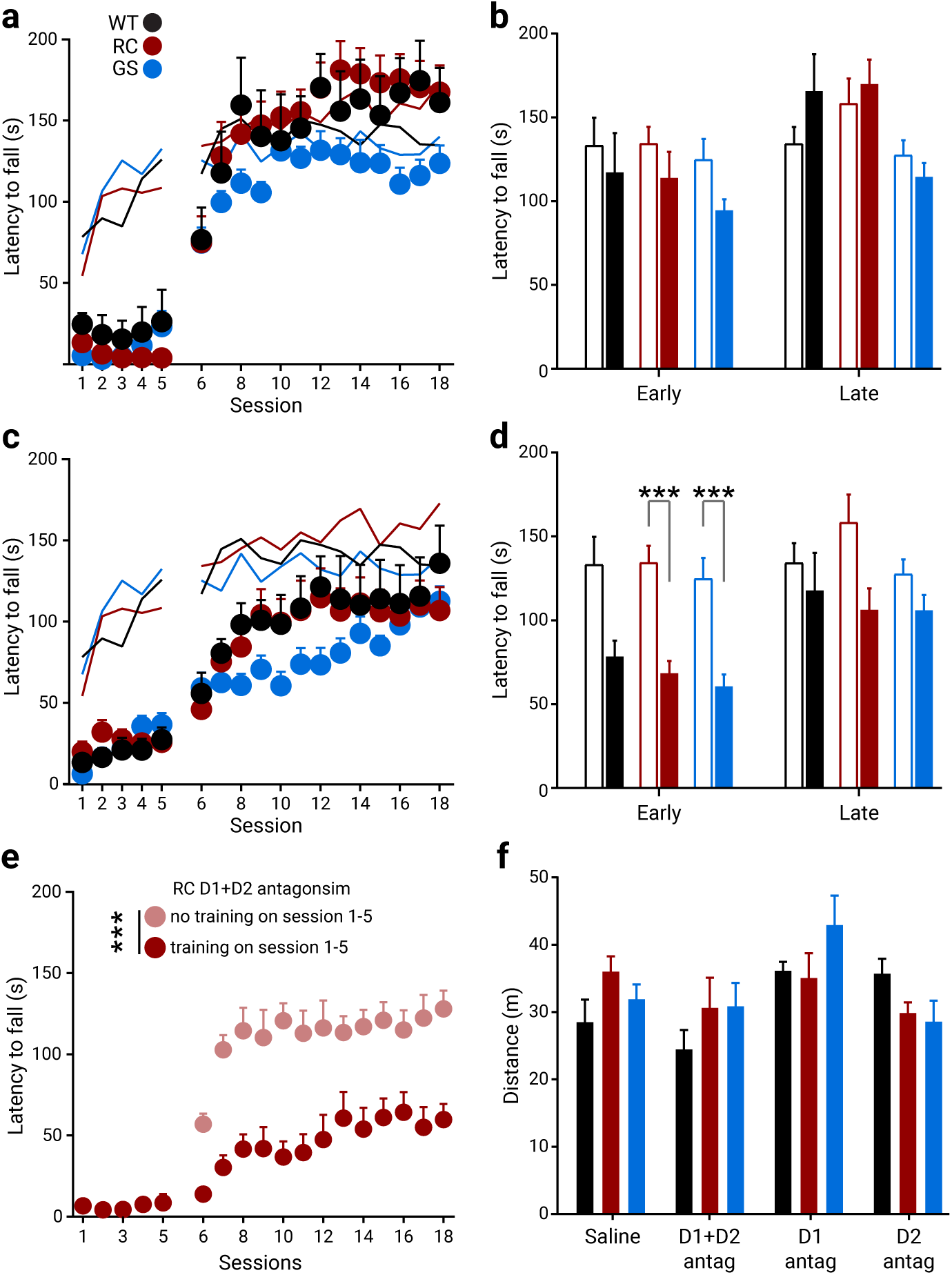
Effect of D1 or D2 antagonism on dopamine-dependent motor learning. Rotarod performance of WT, RC, and GS mice that were administered with an antagonist for either D1 (SCH23390) (**a–b**) or D2 receptors (eticlopride) (**c–d**), 30 min prior to training for the first five days. Solid traces are saline control replotted from Figure 3b for reference. **b** and **d**. Averaged latency of saline control (open bars) and drug-treated mice (filled bars). Early and late refer to session 6–8 and 16–18, respectively, of the no drug recovery phase (D1R antagonist treated groups: n_WT_ = 11, n_RC_ = 11, and n_GS_ = 9 mice; D2R antagonist treated groups: n_WT_ = 9, n_RC_ = 14, and n_GS_ = 10 mice; *** *p <* 0.001, ** *p <* 0.01 vs genotype-matched saline control). **e**. Improved performance of RC mice received antagonist cocktail but no training during the first five sessions. The RC_D1+D2_ group data are from **Figure 3b** for reference (stats). **f**. Distance travelled in the open field after five days of rotarod training with either vehicle or dopamine receptor antagonist cocktail treatment. Another group of mice received dopamine antagonist treatment and rotarod training for five days under the same schedule described in **a–d** and **Figure 3b–c**, and were tested in the open field immediately after the 72-hr break instead of the drug-free phase (saline treated groups: n_WT_ = 11 mice, n_RC_ = 8, and n_GS_ = 7 mice; D1R + D2R antagonist treated groups: n_WT_ = 7, n_RC_ = 6, and n_GS_ = 6 mice; D1R antagonist treated groups: n_WT_ = 7, n_RC_ = 9, and n_GS_ = 9 mice; D2R antagonist treated groups: n_WT_ = 8, n_RC_ = 11, and n_GS_ =7 mice).

**Figure 3−table supplement 1.**
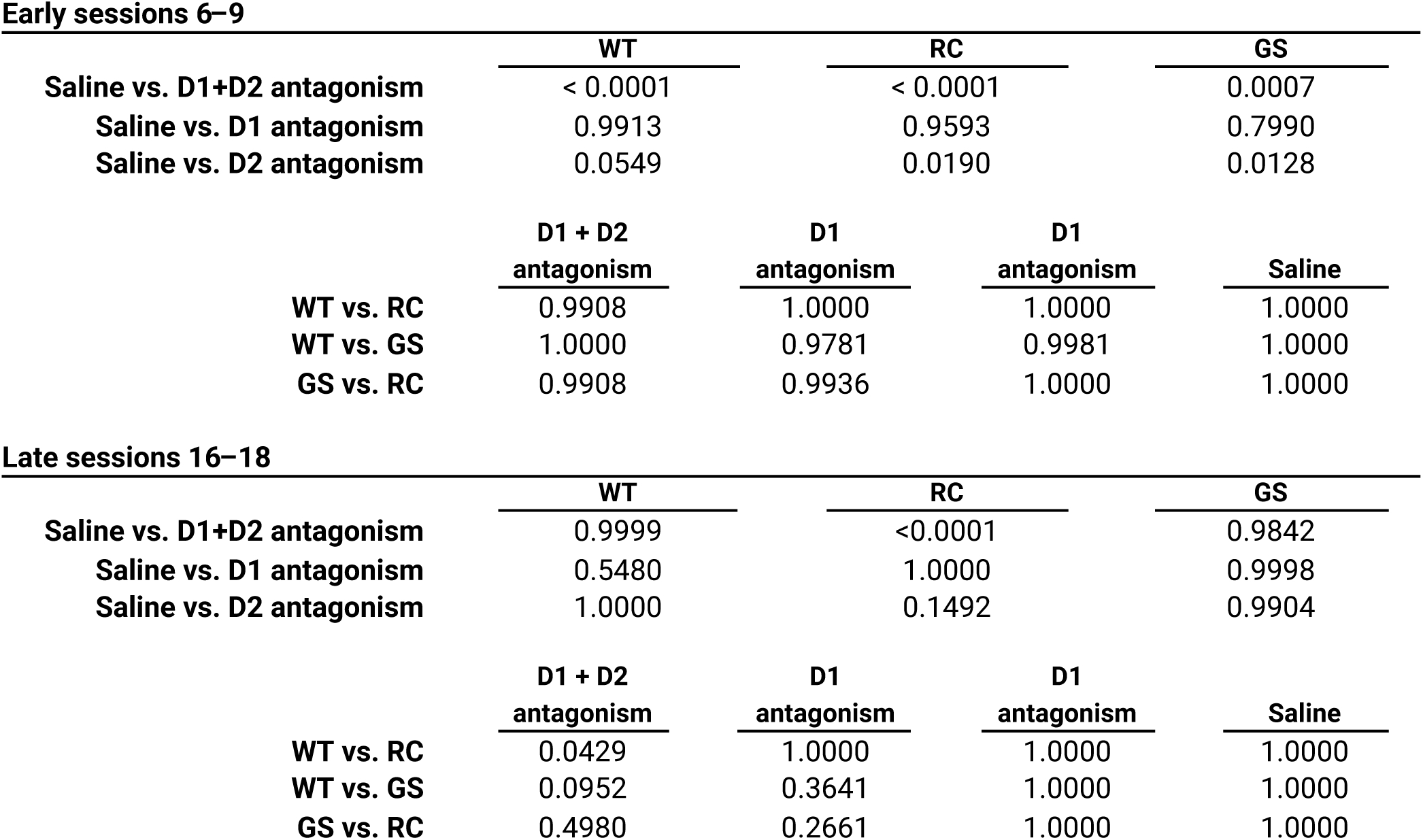
Summary of differences between mutants and pharmacological conditions. Summary table of *p*-values for within-treatment and within-genotype *post hoc* comparisons for both the early and late sessions during the drug-free recovery phase.

